# Forced mouth opening induces post-traumatic hyperalgesia and associated peripheral sensitization after temporomandibular joints injury in mice

**DOI:** 10.1101/2024.01.16.575891

**Authors:** Ishraq Alshanqiti, Hyeonwi Son, John Shannonhouse, Jiaxin Hu, Sinu Kumari, Ghazaal Parastooei, Sheng Wang, Jin Y. Ro, Yu Shin Kim, Man-Kyo Chung

**Affiliations:** Program in Dental Biomedical Sciences, University of Maryland Baltimore, School of Dentistry, Baltimore, MD 21201, USA; Department of Neural and Pain Sciences, School of Dentistry, University of Maryland Baltimore, Baltimore, MD 21201, USA; Department of Basic and Clinical Sciences, School of Dentistry, Umm Al-Qura University, Makkah, 24382, KSA; Department of Oral and Maxillofacial Surgery, School of Dentistry, University of Texas Health Science Center at San Antonio, San Antonio, TX 78229, USA; Programs in Integrated Biomedical Sciences, Translational Sciences, Biomedical Engineering, and Radiological Sciences, University of Texas Health Science Center at San Antonio, San Antonio, TX 78229, USA; Center to Advance Chronic Pain Research, University of Maryland at Baltimore, Baltimore, MD 21201, USA; Program in Neuroscience, University of Maryland at Baltimore, Baltimore, MD 21201; USA

**Keywords:** TMJ, pain, peripheral sensitization, in vivo Ca^2+^ imaging, trigeminal ganglia

## Abstract

Temporomandibular disorder (TMD) is the most prevalent painful condition in the craniofacial area. The pathophysiology of TMD is not fully understood, and it is necessary to understand pathophysiology underlying painful TMD conditions to develop more effective treatment methods. Recent studies suggested that external or intrinsic trauma to TMJ is associated with chronic TMD in patients. Here, we investigated the effects of the TMJ trauma through forced-mouth opening (FMO) in mice to determine pain behaviors and peripheral sensitization of trigeminal nociceptors. FMO increased mechanical hyperalgesia assessed by von Frey test, spontaneous pain-like behaviors assessed by mouse grimace scale, and anxiety-like behaviors assessed by open-field test. In vivo GCaMP Ca^2+^ imaging of intact trigeminal ganglia (TG) showed increased spontaneous Ca^2+^ activity and mechanical hypersensitivity of TG neurons in the FMO compared to the sham group. Ca^2+^ responses evoked by cold, heat, and capsaicin stimuli were also increased. FMO-induced hyperalgesia and neuronal hyperactivities were not sex dependent. TG neurons sensitized following FMO were primarily small to medium-sized nociceptive afferents. Consistently, most TMJ afferents in the TG were small-sized peptidergic neurons expressing calcitonin gene-related peptides, whereas nonpeptidergic TMJ afferents were relatively low. FMO-induced intraneural inflammation in the surrounding tissues of the TMJ indicates potentially novel mechanisms of peripheral sensitization following TMJ injury. These results suggest that the TMJ injury leads to persistent post-traumatic hyperalgesia associated with peripheral sensitization of trigeminal nociceptors.

## 1. INTRODUCTION

Temporomandibular disorder (TMD) refers to a group of musculoskeletal conditions that affect the temporomandibular joint (TMJ), muscles of mastication, and related structures [6]. The incidence of first-onset TMD among U.S. adults is 4% per annum [28]. Based on the Diagnostic Criteria for Temporomandibular Disorders (DC/TMD), TMDs represent multifactorial, heterogeneous, musculoskeletal disorders involving pain and intra-articular structural changes, such as disc displacement and joint degeneration [21]. Therefore, managing patients with TMD is complex, and no one treatment works for all patients. Instead, each patient must be appropriately diagnosed for a specific subtype of TMD and treated via an individually tailored care pathway. Developing more effective treatments should be based on a thorough understanding of the pathogenesis and mechanisms underlying individual etiologic conditions that contribute to painful TMDs. Although diverse pharmacological and nonpharmacological approaches are currently used to manage TMD symptoms [20], effective mechanism-based treatment options are lacking.

Among the diverse etiologies of TMDs, excessive functioning of the TMJ beyond its physiological range can lead to chronic pain, and the annual incidence of TMDs is higher in patients who have experienced a jaw injury [25]. The injury can be intrinsic (i.e., sustained mouth opening and yawning) or extrinsic (i.e., tooth extractions or dental treatments; motor vehicle accidents; accidents resulting in whiplash; oral intubation; sports injuries including falls, bumps, and blows; injuries to the head; and injuries to the neck and shoulder region). The odds ratio of TMD is increased by 3–7 fold in patients who have experienced different types of injuries [24]. To mimic such conditions, animal models have been developed to deliver injuries to the TMJ by adding abnormal TMJ loading or inducing excessive jaw openings in rats and mice [5; 9; 13; 34]. TMJ loading or injury induced by forced mouth opening (FMO) commonly induces facial mechanical hyperalgesia, accompanied by peripheral and central neuroimmune responses, such as activation of macrophages in the trigeminal ganglia (TG) and microglia in the trigeminal nucleus caudalis [34]. As TMJ injuries lead to the degeneration of the condylar cartilage and subchondral bone [11; 13], FMO is a noninvasive, clinically relevant model for post-traumatic hyperalgesia and condylar degeneration. However, pain phenotyping following FMO is primarily limited to assessing mechanical hyperalgesia rather than more clinically relevant outcomes such as jaw function-related pain or emotional disturbances. Furthermore, there is no information regarding FMO-associated functional plasticity of trigeminal afferents. Moreover, sex differences in pain-like behaviors and afferent sensitization in this model have not been determined.

In this study, we investigated clinically relevant pain phenotypes, the neurochemical properties of TMJ afferents, and functional responses of trigeminal afferents following TMJ injury in both male and female mice. A better understanding of post-traumatic TMJ hyperalgesia should improve our ability to investigate the peripheral and central neurobiology of the transition from acute injury to chronic pain and validate the efficacy of newly developed analgesics for treating TMJ hyperalgesia.

## 2. MATERIALS AND METHODS

### 2.1. Experimental animals

We followed the guidelines in the NIH Guide for the Care and Use of Laboratory Animals (Publication 85-23, Revised 1996). The study protocols were approved by the University of Maryland Baltimore or the University of Texas Health Science Center at San Antonio Institutional Animal Care and Use Committee. A 12:12 light-dark cycle was maintained, and all animals had access to food and water ad libitum in a temperature-controlled room. C57BL/6 mice (Jackson Laboratory, Bar Harbor, ME; #000664) or Pirt-GCaMP3 mice [14] were used for the experiments. Adult male or female 8- to 12-week-old mice were used For experiments involving bite force measurements, 16-week-old mice were used. For all behavioral assays, the animals were randomly allocated to experimental groups.

### 2.2. Forced mouth opening (FMO)

The mice were anesthetized with isoflurane inhalation (3% for induction and 1.5% for maintenance). To produce TMJ injury, FMO was performed by placing), a custom-made mouth opener (0.017-inch x 0.025-inch rectangular wire) was between the maxillary and mandibular incisors (Fig. 1A). The mice were kept in the isoflurane chamber for three hours. The FMO procedure was repeated for five days (5D) or two days (2D), as indicated in the result section. The sham group was placed inside the isoflurane chamber without the mouth opener for three hours. A soft diet (DietGel Recovery, ClearH2O) and water gel (HydroGel, ClearH2O) were provided after each FMO procedure. Body weight was closely monitored, and there were no significant changes of body weight occurred between the FMO and sham groups in males and females (Fig. 1B). The mice were randomly allocated to their respective groups, and the experimenters were remained blinded to the group assignment throughout all behavioral assays.

**Figure 1.**
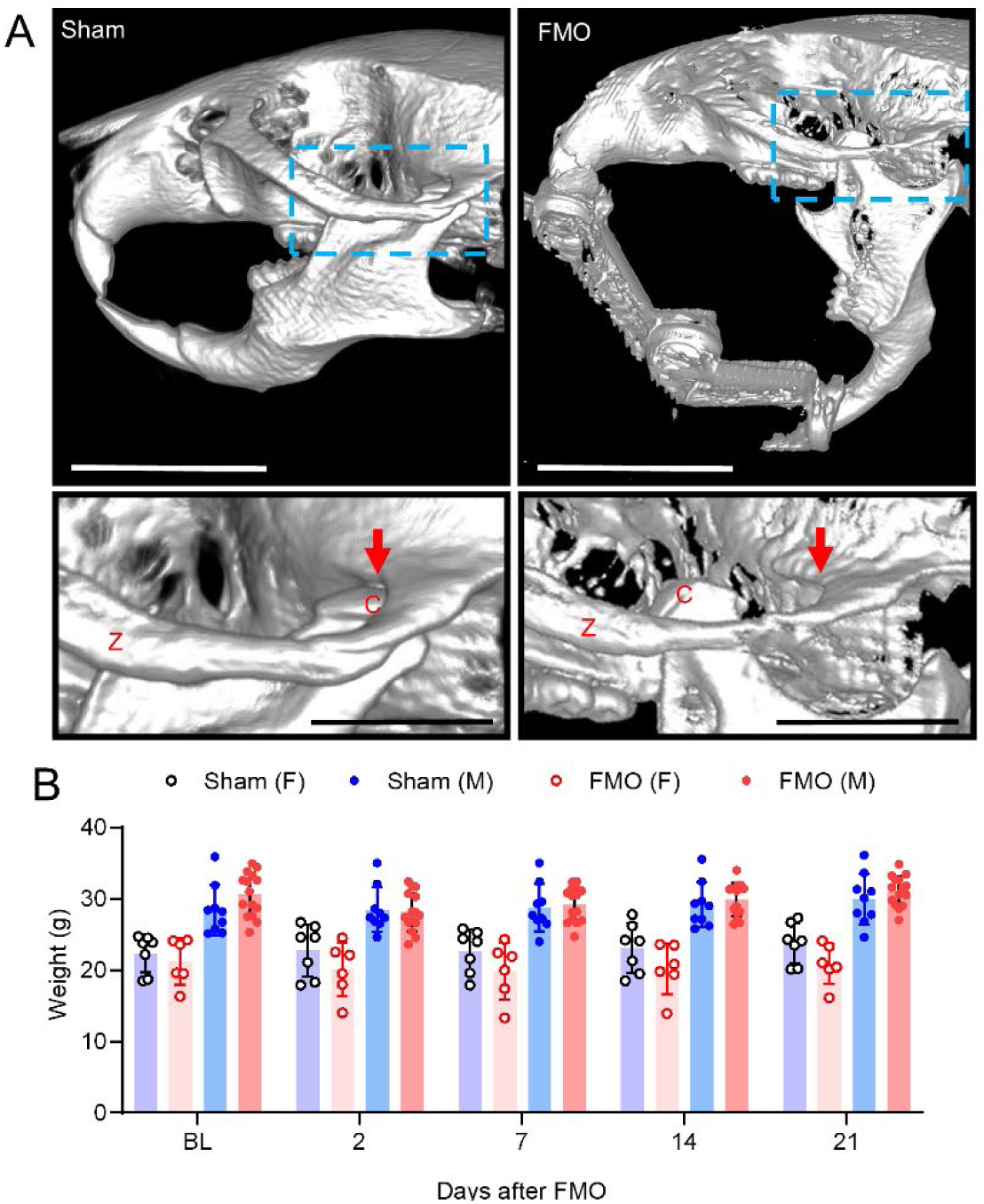
FMO for assessing pain-like behaviors after temporomandibular joint (TMJ) injury in mice. **A.** Micro computed tomography images of two formalin-fixed mice under closed mouth (sham) or FMO conditions. Magnified views of the TMJ areas in the boxes are shown in the panels below. FMO induced movement of the condylar head (c) from the resting position (red arrow) to an extreme anterior position beyond its physiological range and produced excessive stretches, injuries, and inflammation of joint capsules and retrodiscal tissues. FMO, forced mouth opening; z, zygomatic arch; scale bar, 5 mm (top) or 2 mm (bottom). **B.** Body weight of the mice over the course of the experiments. BL, baseline.

### 2.3. Assessment of mechanical sensitivity using von Frey (VF) assay

In order to assess changes in mechanical sensitivity, we conducted the VF assay using up-down techniques (Wang et al. 2020). The orofacial skin over the TMJ was exposed to calibrated VF filaments with bending forces ranging from 0.008 to 4 g. The nocifensive reaction was described as rapid or vigorous head withdrawal from the probing filament. Each VF filament was applied five times, each lasting a few seconds. The response frequencies [(the number of responses/the number of stimuli) × 100%] to a range of VF filament forces were determined, and S-R curves were plotted. We defined the lowest intensity at which animals escaped as the mechanical threshold.

### 2.4. Mouse grimace scale (MGS)

Mouse spontaneous pain can be measured consistently and quantitatively by facial expression coding [35; 36]. Baseline recording followed an acclimatization period of 30 minutes. Two digital video cameras (Sony HDR-CX230/B High Definition Handycam camcorders; Sony Corp., Tokyo, Japan) were positioned to maximize the probability of capturing the faces of the mice. For each experimental time point, mice were videotaped for 30 minutes. Two blinded experimenters extracted the facial images . During video recordings, a clear image of the entire face was manually captured every two to three minutes (10 images per 30-minute session). Three facial action units were scored: orbital tightening, ear position, and whisker change. Each action unit was scored as 0, 1, or 2 based on criteria by blinded experimenters as described previously [35; 36].

### 2.5. Bite force assay

This assay was performed as described previously [36]. To measure bite force, two parallel aluminum plates were connected to a force-displacement transducer (FT03; Grass instruments; Astro-Med Inc., West Warwick, RI, USA) connected with Spike2 software (CED Ltd., Cambridge, United Kingdom). The mice were placed in a modified 60 ml plastic syringe with a wide opening on one side. While manually holding the mouse syringe and moving it at a speed of 0.5–1 cm/sec, the syringe was slowly moved toward the bite plates for the mouse to bite them. To minimize stress, the mouse was immediately released from the syringe if it vigorously moved or attempted to hide inside it. In each session, bite force was measured for 120 seconds, and the top five measurements were averaged. The bite force values measured after FMO or sham procedures were normalized to the baseline obtained from each mouse.

### 2.6. Open Field Test (OFT)

Three days after the final FMO or sham procedure session, each mouse was subjected to open field testing, a well-established and validated test for evaluating exploratory behavior and anxiety in rodents [23]. Briefly, the mice were brought into the testing room and allowed to acclimate for a minimum of 30 minutes before commencing the test. The testing arena, measuring 40 × 40 × 30 cm (length × width × height), was used for the OFT. Each mouse was placed in the center of the arena using their assigned handling method. All testing procedures were conducted between 08:00 and 14:00 hours to minimize the potentially confounding effects of diurnal variations. Mouse activity within the open field arena was tracked for 10 min and recorded using video-tracking software (BehaviorCloud, San Diego, CA, USA). The following parameters were analyzed as validated measures of anxiety: total distance traveled in the central (20 cm X 20 cm) and peripheral zones, time spent in the central zone, central resting and ambulatory time, and central distance traveled.

### 2.7. Labeling of TMJ afferents and immunohistochemistry of the trigeminal ganglia (TG)

Wheat germ agglutinin (WGA) conjugated with Alexa Fluor 488 (WGA-488; Thermo-Fisher Scientific, Invitrogen, Carlsbad, CA, USA) was injected into the TMJ to retrogradely label TMJ afferents. Fur was shaved in the TMJ area, and the location of the TMJ was identified by following the posterior ramus of the mandible and the end of the zygomatic arch. Under ketamine/xylazine anesthesia, WGA-488 (1% in distilled water; 4 µl/joint) was injected into the joint cavity of both TMJs. The next day, the mice were subjected to FMO or sham. Twenty-four hours after completing two days of FMO or sham, the mice were euthanized by transcardial perfusion using 3.7% paraformaldehyde. Immunohistochemical assays of the TG were performed as previously described (Wang et al. 2017; Chung et al. 2011). Tissues were cryoprotected and cryosectioned at 12 μm. Conventional immunohistochemical procedures were performed using rabbit anti-TRPV1 (1:1,000; a generous gift from Dr. Michael Caterina at Johns Hopkins University) and guinea pig anti-calcitonin gene-related peptides (CGRP) (1:1,000; Peninsula Laboratories International, San Carlos, CA, USA)). The sections were incubated with the primary antibodies overnight, followed by secondary antibodies for 60 min. Isolectin B4 (IB4)-conjugated with Alexafluor 568 was labeled for 30 min. We verified the specificity of the primary antibodies using genetically engineered mice lacking the expression of the target gene or by omitting the primary antibody (Chung, Jue, and Dong 2012). We used Image J (NIH, Bethesda, MD, USA) to manually count the immunolabeled neurons and calculate the surface area of the cells. The following criteria for neuronal size classification were used: small < 300 μm^2^, medium 300 to 600 μm^2^, and large > 600 μm^2^ [3; 10]. The experimenters were blinded to the treatment groups during the data analysis step.

### 2.8. Immunohistochemistry of TMJ

Dissected TMJs were decalcified in 0.5M ethylenediaminetetraacetic acid (EDTA) in PBS (pH 7.4) for two weeks at room temperature for 2 weeks at room temperature. The decalcified tissues were dehydrated, embedded in paraffin, and sectioned into 5 µm slices. Immunohistochemical assays were performed using anti-CD45 (Abcam, Cambridge, United Kingdom; #ab10558; rabbit). The sections were stained with 4’,6-diamidino-2-phenylindole (DAPI) to visualize the cellular nuclei.

### 2.9. Micro-focus computed tomography

Micro-focus computed tomography (µCT) images of the skull were obtained using a Siemens Inveon Micro-PET/SPECT/CT (Siemens, Ann Arbor, MI, USA). Siemens Inveon Research Workplace 4.2 software was used for image acquisition, processing, and 3-dimensional image viewing.

### 2.10. In vivo GCaMP calcium imaging

To determine the effects of the FMO on the transduction of noxious stimuli at nociceptive nerve terminals, we performed in vivo Ca^2+^ imaging of the TG of anesthetized mice expressing GCaMP3 from the Pirt locus [14]. Mice were anesthetized via i.p. administration of a ketamine/xylazine mix of 80/10 mg/kg (MilliporeSigma, St Louis, MO, USA). An ophthalmic ointment (Lacri-lube; Allergen Pharmaceuticals, Dublin, Ireland) was applied to the eyes for protection. The dorsolateral portion of the skull on the right side was exposed by removing the skin and muscle. Using a dental drill (Buffalo Dental Manufacturing, Syosset, NY, USA), the parietal bone between the right eye and ear was excised to create a cranial window (∼10 X 10 mm). The TG was subsequently visualized by gently aspiring the cortical tissues above it. Each mouse’s body temperature was maintained throughout the surgery by a heating pad and rectal probe at 37 °C ± 0.5 °C.

In vivo, intact TG *Pirt*-GCaMP3 Ca^2+^ imaging in live mice was performed for two to five hours immediately after exposure surgery, as previously described (Kim et al., 2016). Following exposure surgery, each mouse was positioned using a custom-designed platform under a microscope. To minimize any potential disruptions caused by respiratory and cardiac activities, each mouse’s head was fixed by a head holder. During imaging, body temperature was maintained at 37 °C ± 0.5 °C, with anesthesia sustained at 1–2% isoflurane dispensed from a vaporizer using pure oxygen. Live images were captured in ten frames per cycle in frame-scan mode at ∼4.5 to 8.79 s/frame, ranging from 0 to 90 µm, using a 5 X 0.25 NA dry objective at 512 X 512 pixels or higher resolution with solid diode lasers tuned at 488 nm and emission at 500–550 nm. External stimuli, including VF filaments (0.4 g) and noxious cold (4 °C) or hot water (50 °C), were sequentially administered to facial skin in the mandibular region. Additionally, capsaicin (500 mM, 10 μl) was injected intracutaneously into the different TG branches. To avoid unwanted mechanical stimulation during the injection process, we inserted a needle into mandibular skin prior to the initiation of the recording.

Raw image stacks were collected, deconvoluted, and imported into ImageJ (NIH) for imaging data analysis. Consecutive optical planes were re-aligned and corrected using an image alignment plugin based on stackreg rigid-body cross-correlation. Ca^2+^ signal intensity was represented as the ratio of F_t_ (fluorescence intensity in each frame) to F_0_ (mean fluorescence intensity during the first one to four frames). All responding cells were verified by manually inspecting the raw imaging data. The size of TG neurons was classified as small (<20 µm), medium (20-25 µm), and large (>25 µm) neurons according to the diameter.

### 2.11. Statistical Analysis

Data are presented as mean ± standard error of the mean. Statistical comparisons were performed using Student’s t-tests or an analysis of variance (ANOVA) followed by a post hoc test as indicated in the figure legends. A p-value < 0.05 was considered statistically significant. All statistical analyses were performed using Prism (GraphPad Software, La Jolla, CA, USA)

## 3. RESULTS

### 3.1. FMO induced long-lasting increases in pain-like behaviors and anxiety-like behaviors

Results of the VF assay on mandibular skin showed that the head withdrawal threshold was significantly lower in the FMO group than the sham group for nine days following the completion of FMO 5D (Fig. 2A). There was no sex difference in FMO-induced mechanical hyperalgesia (Fig. 2B). Mechanical hyperalgesia was robust until 21 days following FMO 5D (Fig. 2C), suggesting that FMO induced long-lasting mechanical hyperalgesia in the face. In contrast, mechanical hyperalgesia produced by FMO 2D was recovered after 18 days, suggesting that the duration of mechanical hyperalgesia depends on the extent of TMJ injury.

**Fig. 2.**
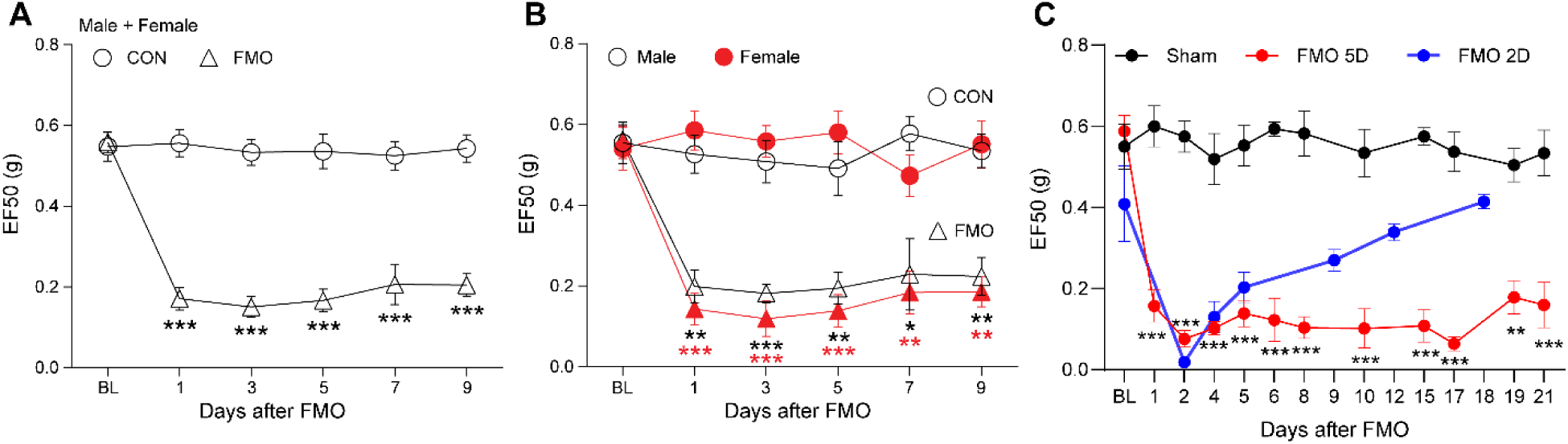
FMO induces long-lasting mechanical hyperalgesia of the skin in the V3 region. Withdrawal thresholds (EF50) tested assessed using the von Frey filament after FMO 5 days (5D) are shown for all mice (**A**; n = 12 mice per group) and a comparison between males and females (**B**,**B**; n = 6 mice per group). In a separate cohort, the von Frey assay was performed over the long-term (**C**; n = 6 mice per group) after either FMO for five days (FMO 5D) or for two days (FMO 2D). Circles correspond to the CON group, and triangles represent the FMO group. CON: control, FMO: forced mouth opening. Error bars indicate S.E.M. *p < 0.05; **p < 0.01; ***p < 0.001 in Sidak’s post hoc test (**A** and **C**) or Turkey’s post hoc test (**B**) following a two-way repeated measure ANOVA.

MGS (a measure of spontaneous pain-like behaviors) scores were elevated after one or three days following the initiation of the first FMO session (Fig. 3A). MGS peaked two days after the completion of FMO 5D and was slightly reduced thereafter. The extent of the MGS increase on day 21 was modest but still significantly higher than in the sham group. Changes in MGS were not significantly different between males and females (Fig. 3B).

**Figure 3.**
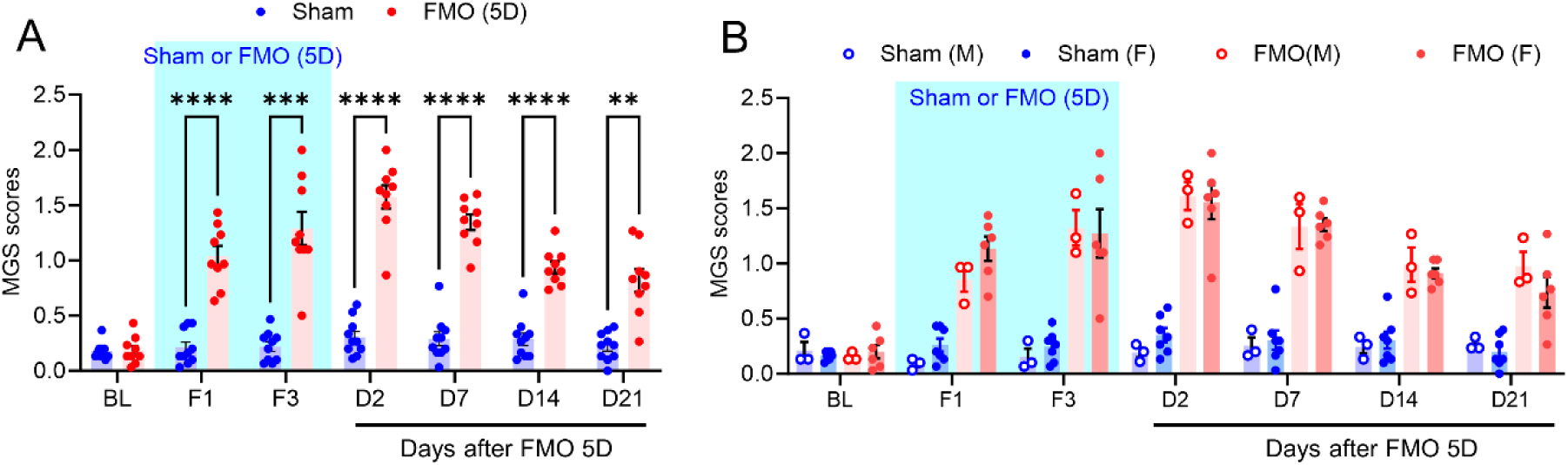
FMO induced long-lasting spontaneous pain-like behaviors. **A.** The MGS was assessed at BL during the 5-day period of FMO (F1, F3), and after the completion of FMO 5D (D2, D7, D14, and D21). N=10 mice in the sham and n=9 in the FMO group. **p < 0.01; ***p < 0.001; ****p<0.0001 using the Bonferroni post hoc test following a two-way ANOVA. FMO, forced mouth opening; MGS, mouse grimace scale; BL, baseline. **B.** Comparison of MGS in males (n=3 per group) and females (n=7 in sham and n=6 in FMO).

The BF assay was conducted to evaluate how FMO affected jaw functioning. We found that mice that underwent FMO 5D were discouraged from biting and exhibited highly inconsistent performances, which made evaluating the assay difficult. Therefore, we performed FMO for only 2 days (FMO 2D) for the BF assay. After FMO 2D, less than 50% of the mice performed well exhibited successful biting (Fig. 4A). The proportion with a successful bite increased over time, which was not significantly different between males and females, and recovered close to the level of the sham group 19 days after FMO 2D. When we compared the bite force of FMO mice with successful bites to sham mice, it was significantly decreased in the FMO group compared to the sham until day 19 following FMO 2D (Fig. 4B). There was no difference between male and female mice (Fig. 4C and D).

**Fig. 4.**
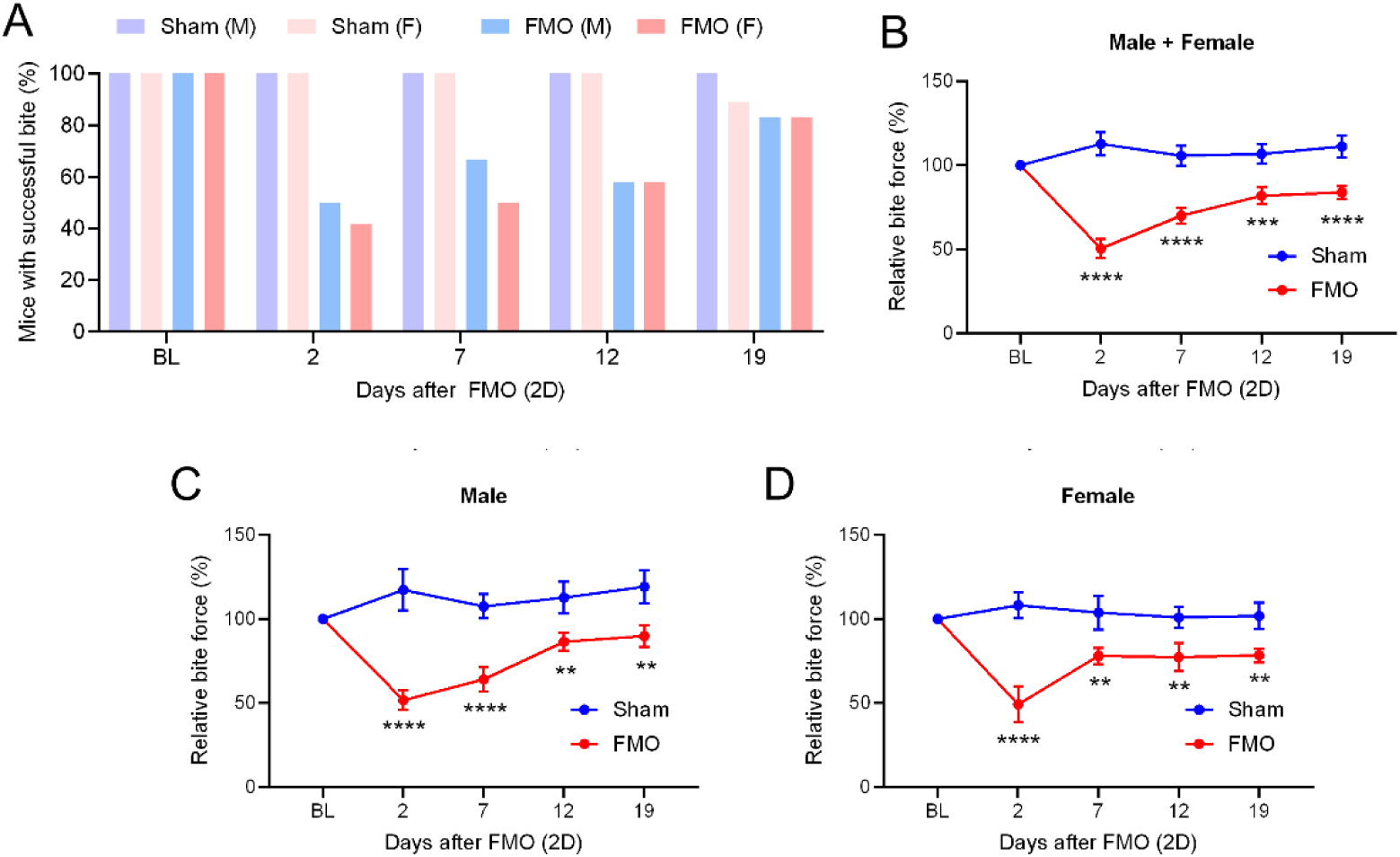
FMO induced prolonged reduction in bite force. **A.** The proportion of mice showing successful bites in the bite force assay in males and females at BL and throughout the experimental period after two days of FMO or sham. Data were taken from 19 male and 19 female mice (7 in the sham and 12 in FMO groups). **B-D.** Normalized bite force in which force values measured after FMO or sham were normalized to their respective BL values in all (**B**), male (**C**), and female mice (**D**). N=14 (sham) and 24 (FMO) in **B**. N=7 (sham) and 12 (FMO) in **C** and **D**. *p<0.05, **p<0.005, ***p<0.0005, ****p<0.0001 in mixed-effects analysis followed by Sidak multiple comparisons between sham and FMO mice. FMO, forced mouth opening; BL, baseline; 2D, two days.

We also assessed anxiety-like behaviors after FMO using OFT. FMO and sham mice exhibited comparable total distance traveled and ambulatory time (Fig. 5A-C). However, the FMO mice showed significantly decreased total time spent in the central area and central resting time (Fig. 5D-E).

**Figure 5.**
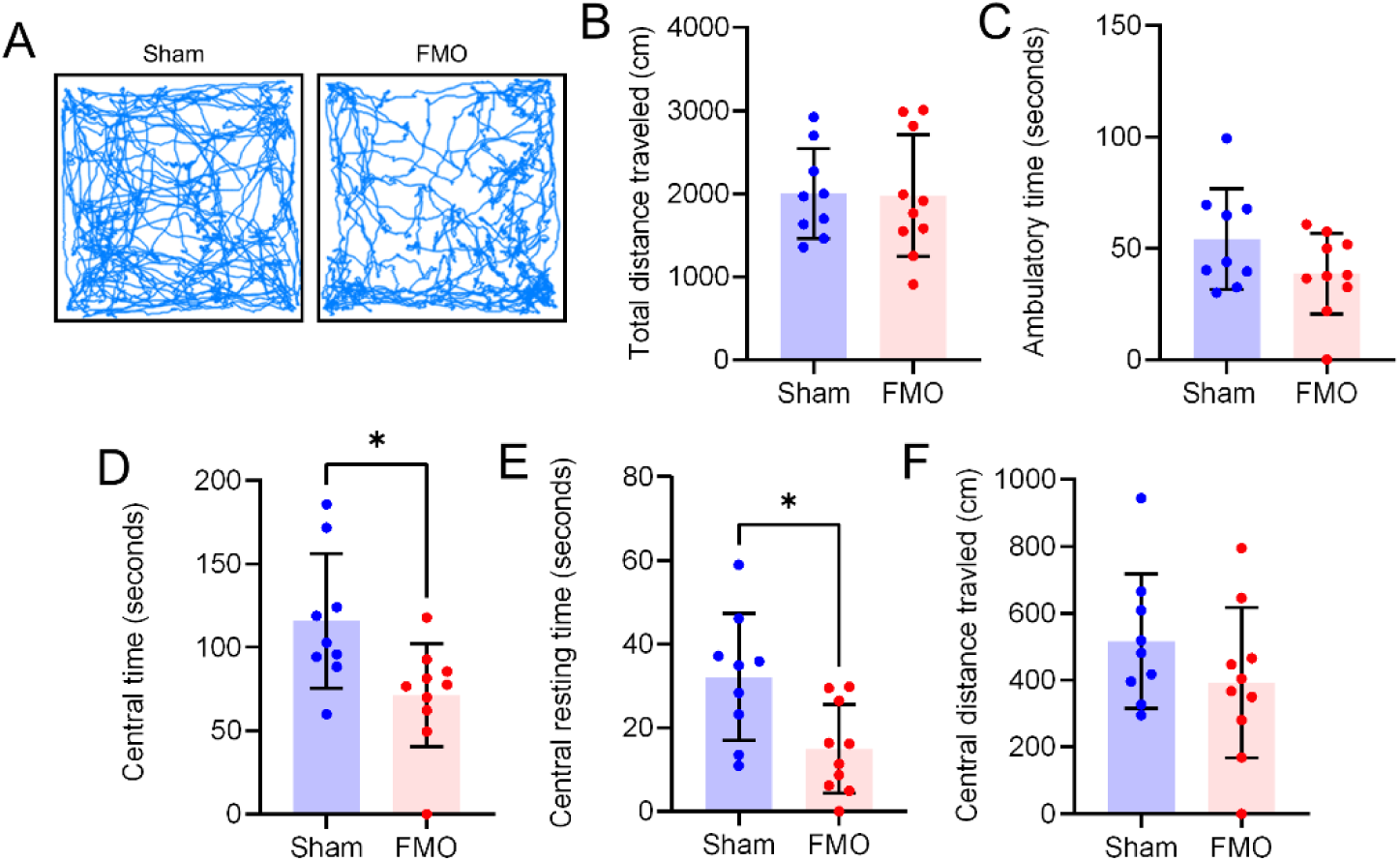
FMO induced anxiety-like behaviors in mice. **A.** Representative traces of mouse movements in a box during 10 minutes of video recording 2days after sham or FMO 5days. **B-F**. Comparison of the total distance traveled (**B**), ambulatory time in the central zone (**C**), travel time in the central zone (**D**), resting time in the central zone (**E**), and distance traveled in the central zone (**F**). *p<0.05 using Student’s unpaired t-test. N=9 in sham and 10 in FMO groups. FMO, forced mouth opening.

### 3.2. FMO induced hypersensitivity of TG neurons

We monitored the activity of TG neurons in sham and FMO mice using in vivo intact TG Pirt-GCaMP3 Ca^2+^ imaging. During the imaging, the TG neurons were spontaneously activated without stimulus, which was not confined to the V3 region of the TG but was also observed in the ophthalmic (V1) and maxillary (V2) areas (Fig. 6A). The spontaneous activities were greater in the FMO than the sham group (Fig. 6B). The amplitudes of Ca^2+^ transients assessed by the area under the curve (AUC) were significantly greater in the FMO than the sham group (Fig. 6C). The total number of spontaneously activated neurons was significantly increased in FMO mice compared to sham (Fig. 6D). The increases in spontaneous activities in FMO mice were primarily occurred in small (<20 µm)- and medium-diameter (20-25 µm) neurons. In both the control and FMO groups, we found no statistically significant difference between males and females (Fig. 6E).

**Figure 6.**
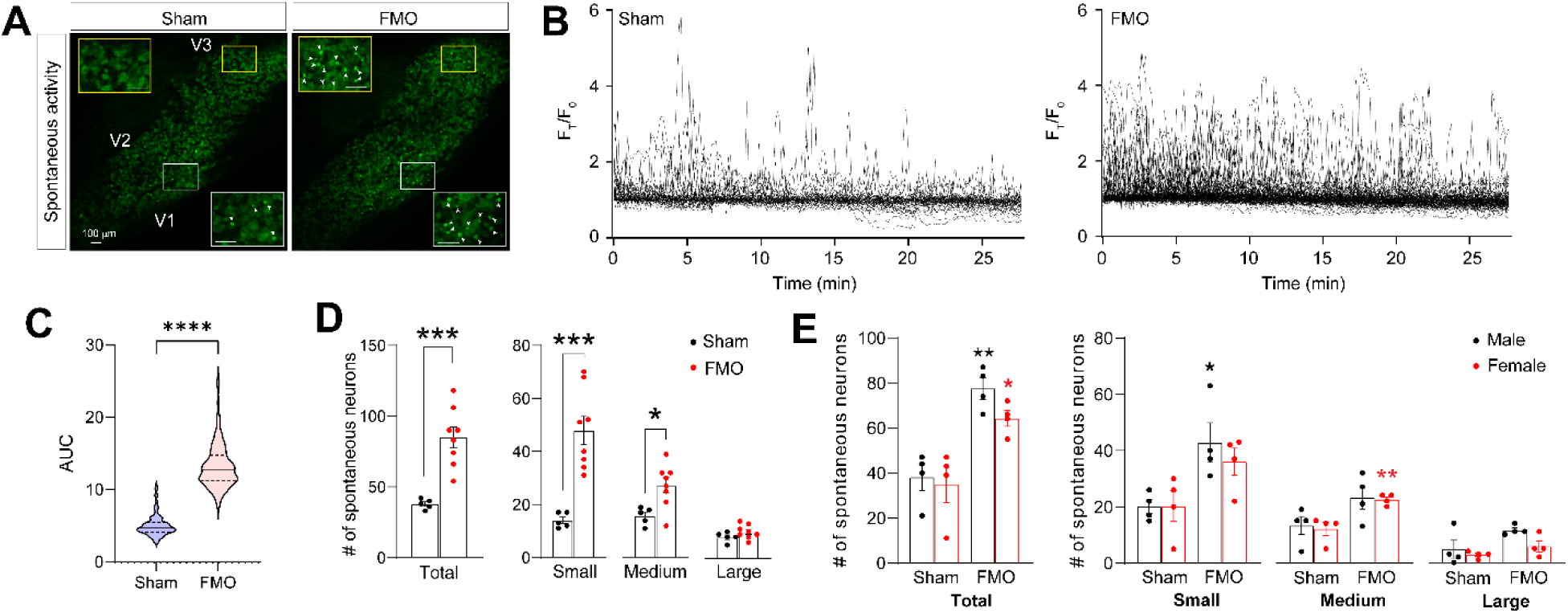
FMO increased spontaneously activated neurons in the intact TG using in vivo Pirt-GCaMP Ca^2+^ imaging. **A.** Representative images of neurons showing spontaneous activities in intact TG *Pirt*-GCaMP3 Ca^2+^ in vivo imaging one or two days after FMO 5D. V1 (ophthalmic), V2 (maxillary), and V3 (mandibular) areas within the TG indicate the location of neuronal cell bodies in TG imaging. Magnified images of the indicated areas are shown in the insets. Scale bars, 100 µm. White arrowheads indicate activated neurons. **B**. Representative superimposed normalized Ca^2+^ transient traces in TG neurons from mice in the sham (left) and the FMO group (right). **C.** Area under the curve (AUC) of normalized Ca^2+^ transients between the sham and FMO group. Solid line within the plot, median; dotted lines, quartiles. N=200 neurons per group. ****p<0.0001 in in Mann-Whitney test. **D.** The total number of spontaneously activated neurons and according to the cell diameter of each group in control and FMO mice. *p < 0.05; ***p < 0.001 using the unpaired Student’s t-test. N=5 in sham and n=8 in the FMO groups. **E.** A comparison of the number of spontaneously activated neurons per TG in total or in different cell diameter groups in Sham and FMO mice based on sex (black for male and red for female). N=4 mice per group. *p < 0.05; **p < 0.01 (sham vs FMO) using Turkey’s post hoc test following a two-way ANOVA. There was no statistical significance between male and female.

To assess responses of TG neurons to mechanical stimuli, we applied 0.4-g VF filaments to the skin region innervated by mandibular division (V3) of trigeminal nerves. In the V3 regions of TG, mechanically activated neurons were more abundant in the FMO than the sham (Fig. 7A). The Ca^2+^ transient changes were greater in the FMO than the sham (Fig. 7B-C). The total number of spontaneously activated neurons was significantly increased in FMO mice compared to sham (Fig. 7D). The increases in mechanical activation in FMO mice were primarily due to the increases in the activation of small-diameter neurons. In both the control and FMO groups, we found no statistically significant difference between males and females (Fig. 7E).

**Figure 7.**
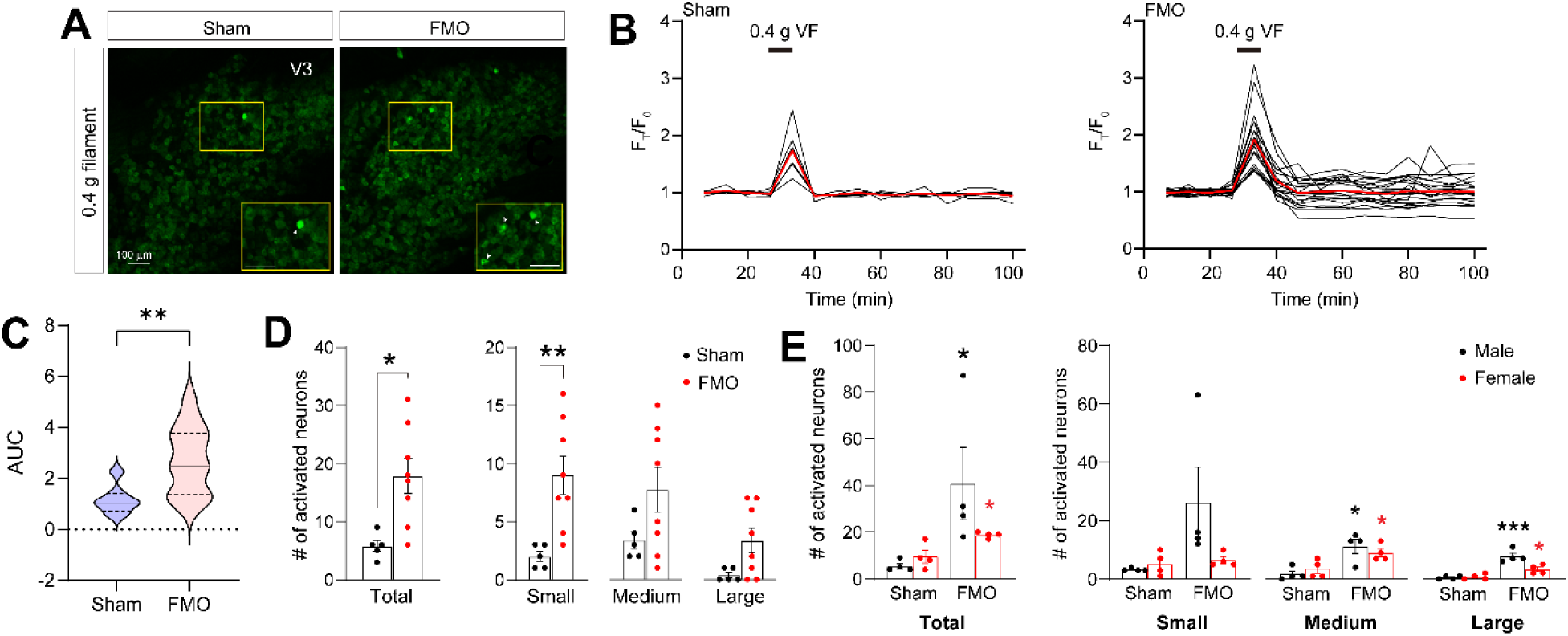
FMO increased TG neurons activated by mechanical stimuli on facial skin in vivo Pirt-GCaMP Ca^2+^ imaging. **A.** Representative images showing mechanically activated neurons in intact TG *Pirt*-GCaMP3 Ca^2+^ in vivo imaging one or two days after FMO 5D. V3 (mandibular) areas within the TG indicate the location of neuronal cell bodies in TG imaging. Magnified images of the indicated areas are shown in the insets. Scale bars, 100 µm. White arrowheads indicate activated neurons. **B**. Representative superimposed normalized Ca^2+^ transient traces in TG neurons from mice with 0.4-g von frey (VF) stimulation on the mandibular skin in the sham (left) and the FMO group (right). The red traces represent the averaged traces in each group. **C.** Area under the curve (AUC) of normalized Ca^2+^ transients between the sham and FMO group. Solid line within the plot, median; dotted lines, quartiles. N=7 neurons in sham and 21 neurons in FMO. **p<0.01 in in Student’s t-test. **D.** The total number of mechanically activated neurons and according to the cell diameter of each group in control and FMO mice. *p < 0.05; **p < 0.01 using the unpaired Student’s t-test. N=5 in sham and n=8 in the FMO groups. **E.** A comparison of the number of mechanically activated neurons per TG in total or in different cell diameter groups in Sham and FMO mice based on sex (black for male and red for female). N=4 mice per group. *p < 0.05 (sham vs FMO) using Turkey’s post hoc test following a two-way ANOVA. There was no statistical significance between male and female.

We also assessed the responses of TG neurons to cold, hot, and capsaicin stimuli on V3 skin (Fig. 8). When cold water (4°C) was applied to the V3 region, the activated TG neurons were significantly higher in FMO mice compared to sham, and small-diameter TG neurons were significantly increased in FMO mice (Fig. 8A,B). Activations of neurons by hot water (50°C) applied to skin in the V3 region were not significantly different between FMO and sham mice (Fig. 8C,D). The number of TG neurons activated by capsaicin injection into the V3 region were significantly greater in FMO than sham mice, particularly medium-diameter ones (Fig. 8E,F). In both the control and FMO groups, we found no statistically significant difference between males and females (Fig. 8E). Overall, these data suggest that FMO induced hypersensitivity of TG neurons without sex differences.

**Figure 8.**
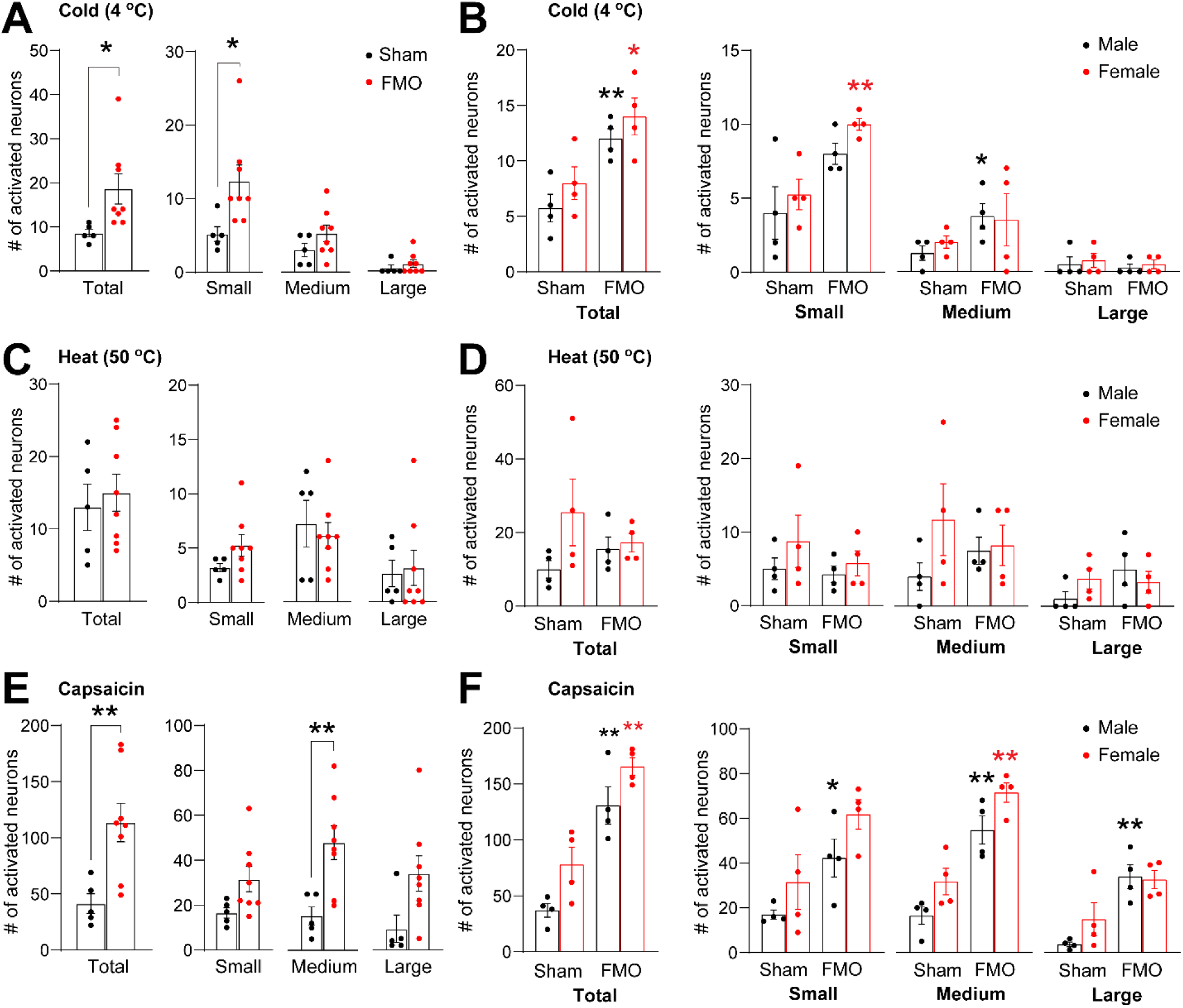
FMO increased cold-and capsaicin-activated, but not heat-activated, neurons in the intact TG using in vivo Pirt-GCaMP Ca^2+^ imaging. **A, C, E.** The total number of TG neurons activated by cold stimuli (4 °C water; **A**), heat stimuli (50 °C water; **C**), or capsaicin injection (E). *p < 0.05; **p < 0.01 using the unpaired Student’s t-test. N=5 in sham and n=8 in the FMO groups. **B, D, F.** A comparison of the number of mechanically activated neurons per TG in total or in different cell diameter groups in sham and FMO mice based on sex (black for male and red for female). N=4 mice per group. *p < 0.05; **p < 0.01 (sham vs FMO) using Turkey’s post hoc test following a two-way ANOVA. There was no statistical significance between male and female.

### 3.3. FMO induced neuroinflammation in the TMJ

We examined the neurochemical properties of TMJ afferents in the FMO and sham groups. Following the TMJ injection of the retrograde labeling dye WGA-488, FMO or sham procedures were performed, and immunohistochemical assays were conducted to determine changes in the neurochemical properties of TMJ afferents (Fig. 9A). The majority (159 out of 308; 53%) of WGA+ neurons in the sham group were small neurons (<300 µm^2^). The proportion of small neurons was slightly higher in FMO mice (256 out of 411; 63%) (Fig. 9B). Consequently, the overall size of the WGA+ neurons in the FMO group was significantly smaller than the sham group (Fig. 9C). The majority of WGA+ neurons were CGRP+ (Fig. 9D). Approximately 20% of WGA+ TMJ afferents expressed TRPV1, whereas there was fewer IB4-labeled WGA+ neurons (Fig. 9E-F). The neurochemical properties of WGA+ TMJ afferents were not significantly different between the sham and the FMO groups (Fig. 9D-E).

**Figure 9.**
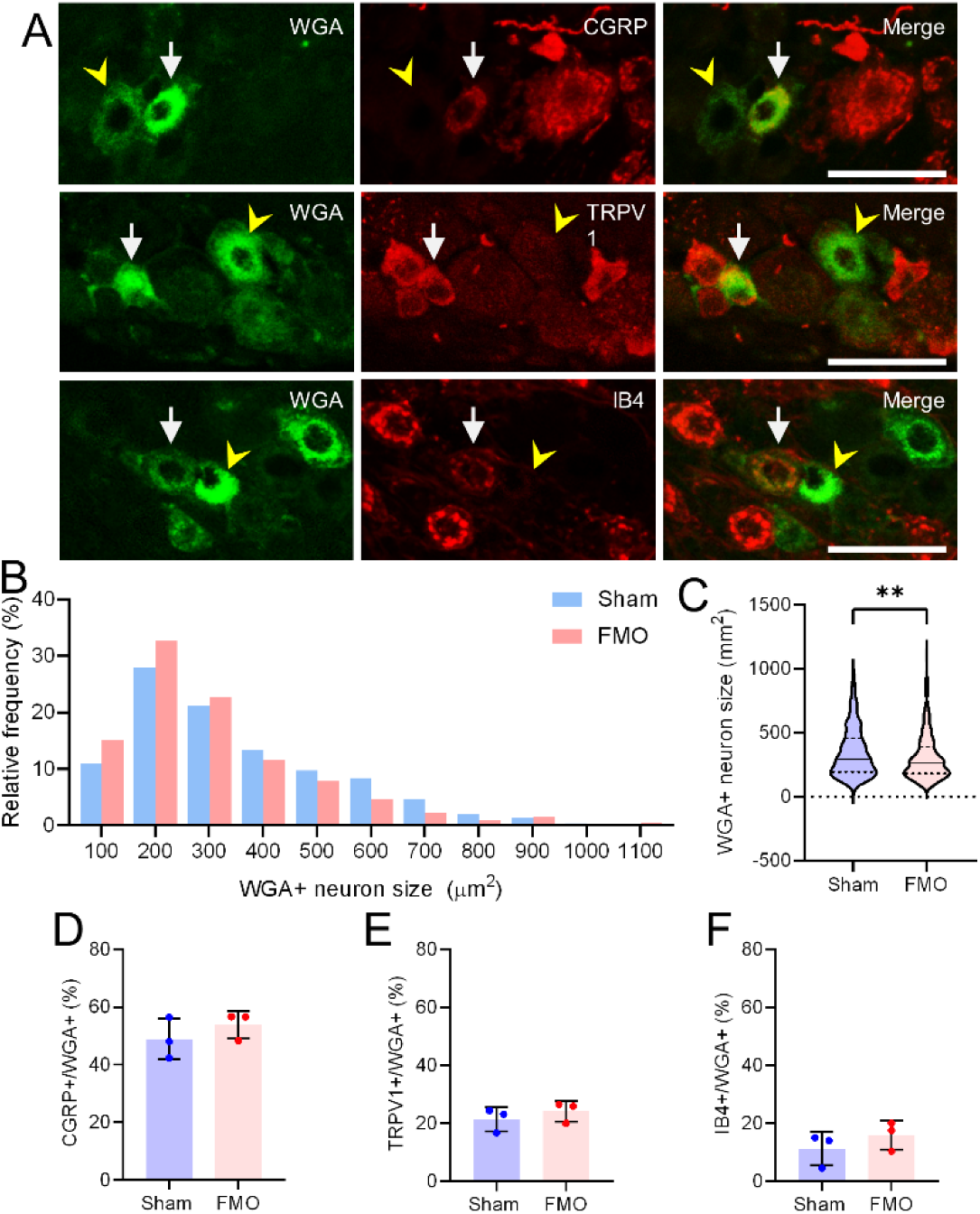
Retrograde-labeled TMJ afferents in mice TG. **A.** Representative images of WGA-labeled TMJ afferents co-localized with CGRP, TRPV1, or IB4 as indicated. Arrows show examples of WGA+ neurons co-localized with the indicated markers. Arrow heads show examples of WGA+ neurons that are not co-localized with the indicated markers. TMJ, temporomandibular joint; WGA, wheat germ agglutinin. Scale, 50 µm. **B.** Size distribution of WGA+ neurons in the TG of mice in sham and FMO groups. TG, trigeminal ganglia. **C.** Comparison of the size of WGA+ neurons. **P<0.01 using the Student’s t-test. N=307 in sham and 407 in FMO groups. **D-F**. Proportion of CGRP+ (**D**), TRPV1+ (**E**), and IB4+ (**F**) neurons among WGA+ neurons. N=3 TG in each group. Total numbers of WGA+ neurons analyzed in each group were 303 (sham) and 413 (FMO) in panel **D**, 219 (sham) and 165 (FMO) in panel **E**, and 389 (sham) and 295 (FMO) in panel **F**.

In decalcified TMJ tissues, we found dense infiltration of CD45+ inflammatory cells in the connective tissues surrounding the TMJ after FMO (Fig. 10A). In sagittal sections of the TMJ, we observed the cross-section of nerve bundles in which DAPI+ cells were located. The density of intraneural DAPI+ cells was significantly higher in the FMO group than in the sham group (Fig. 10B-D). Likewise, CD45+ immune cells were significantly greater in the nerve bundles of the FMO mice than in the sham mice (Fig. 10E).

**Figure 10.**
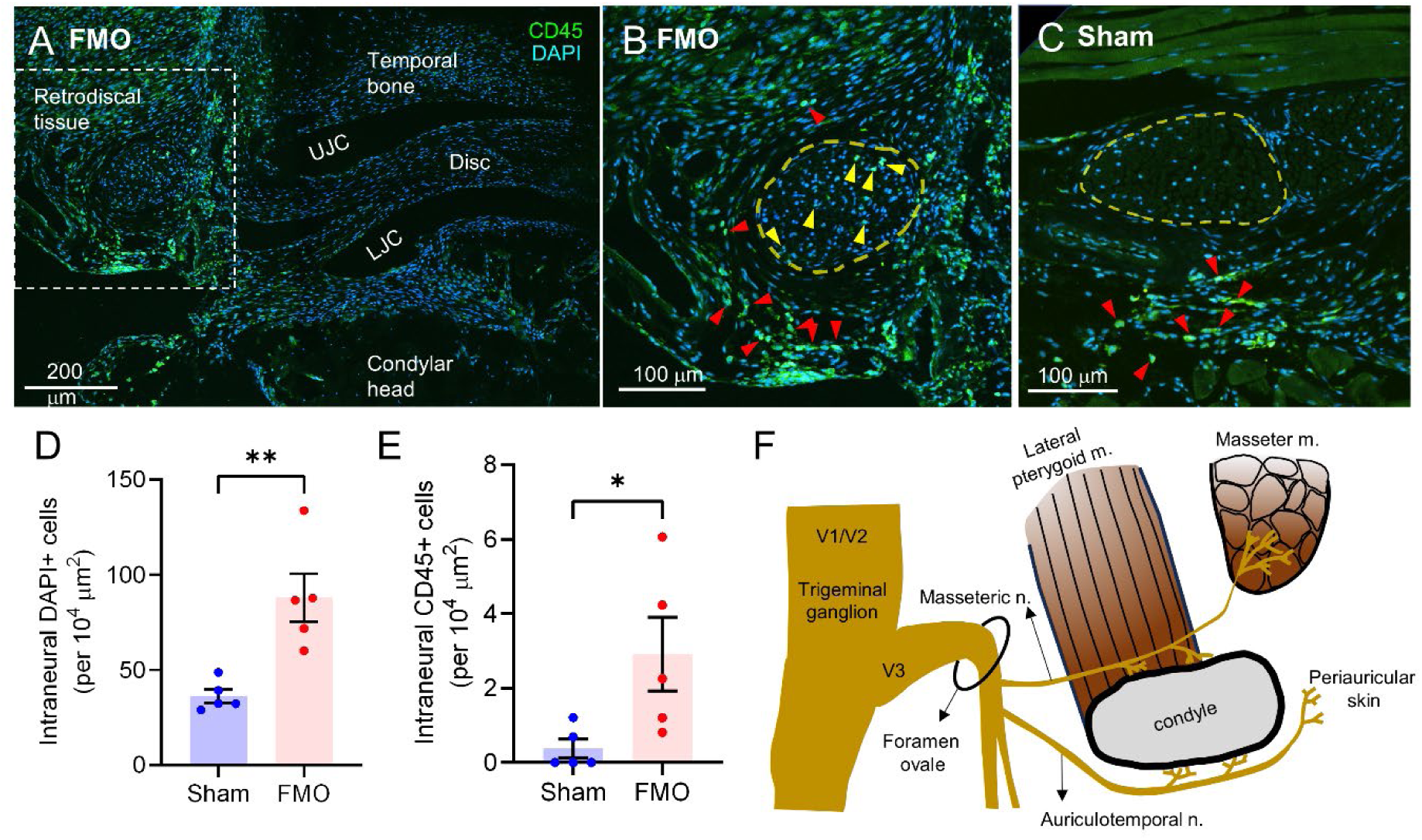
Intraneural inflammation in the surrounding area of the TMJ after FMO. **A**. Sagittal sections of the TMJ three days following FMO 5D after immunohistochemical labeling with CD45, a marker of nucleated immune cells, (green) and DAPI. TMJ, temporomandibular joint; FMO, forced mouth opening; 5d, 5 days; LJC, lower joint cavity; UJC, upper joint cavity. **B-C.** Cross section of the nerve bundle in the retrodiscal area of the TMJ in the FMO (**B**) or sham (**C**) group. The image in **B** is a magnification of the indicated areas in **A**. Red arrow heads indicate examples of CD45+ cells in the tissue. The dotted lines in **B** and **C** represent a nerve bundle. Yellow arrow heads indicate CD45+ cells inside the nerve bundle. **D-E.** Quantification of intraneural DAPI+ cells (**D**) or CD45+ cells (**E**) per unit area. *p<0.05 and **p<0.01 using Student’s t-test. N=5 mice per group. **F.** Diagram of the horizontal view of the human TMJ and its innervation. The masseteric and auriculotemporal nerves are shown coursing laterally, passing the TMJ anteriorly and posteriorly, respectively, and innervating the TMJ, masseter muscles, and the periauricular skin.

## 4. DISCUSSION

We found that TMJ injury produced long-lasting mechanical hyperalgesia of the face, spontaneous pain-like behaviors, disturbances of jaw functions, and anxiety-like behaviors—all of which mimic the clinical manifestation of TMJ arthralgia. Therefore, along with the previous studies [5; 9; 13; 34], our results suggest that FMO-induced TMJ injury in mice is a well-justified, noninvasive model for simulating clinically relevant phenotypes of persistent pain and peripheral sensitization in painful conditions such as TMJ arthralgia and TMJ arthritis.

In our *in vivo* GCaMP Ca^2+^ imaging, we found the number of TG neurons activated spontaneously or by noxious stimuli was increased in the FMO compared to the sham group. This was accompanied by the increased amplitudes of the Ca^2+^ transients. The increased afferents were mostly small to medium-sized neurons, suggesting sensitization of nociceptors. These results indicate that FMO induced peripheral sensitization of the trigeminal nociceptors. Consistently, most back-labeled TMJ afferents are small- to medium-sized neurons. Our immunohistochemical assays in the TG also showed that most of these responses are likely mediated by peptidergic nociceptors expressing CGRP. Innervation of CGRP-expressing nerve terminals within the human TMJ has been reported [7]. The contribution of CGRP to hyperalgesia following TMJ inflammation or masseter muscle tendon ligation in mice has also been demonstrated [27; 33]. FMO did not increase the proportions of CGRP or TRPV1-expressing TMJ afferents in the TG, which is inconsistent with the increased capsaicin sensitivity of TG neurons following FMO. We assume that the capsaicin hypersensitivity is attributable to post-translational modification, such as phosphorylation of TRPV1 [12], rather than ectopic expression of TRPV1 in the TG. More detailed neurochemical and genetic changes within the TG following FMO need to be investigated in the future.

Our behavioral assays and in vivo imaging collectively support the idea that peripheral sensitization is associated with behavioral hyperalgesia following TMJ injury. TMJ injury led to increased spontaneous pain-like behaviors, consistent with increased spontaneous firing of TG neurons. TMJ injury also increased mechanical hyperalgesia on the face, consistent with increased mechanical sensitivity of TG neurons. Peripheral sensitization was not sex-dependent, consistent with the lack of sex difference in hyperalgesia. These results support the idea that peripheral nociceptor sensitization primarily drives the observed hyperalgesia. Interestingly, neuronal responses to noxious heat applied to the facial skin were not significantly different between sham and FMO groups, suggesting no significant thermal hypersensitivity of primary afferents developed after FMO. Although we were unable to determine the thermal sensitivity of facial skin in mice, a recent analysis using an operant orofacial pain assessment device (OPAD) showed that an intra-TMJ injection of carrageenan produced modest thermal hyperalgesia on facial skin on the first day but not afterward [17]. In contrast, cold hypersensitivity of TG afferents was evident after FMO in our TG recordings, which is consistent with the cold hyperalgesia observed in the data from the OPAD assay in rats [17].

Although FMO produces TMJ injury, we do not interpret all nocifensive or neuronal responses as being derived exclusively from the TMJ. Increased spontaneous pain-like behaviors, decreased bite force, or increased anxiety-like behaviors were likely due to the concerted activations of nociceptors in the TMJ and adjacent orofacial tissues, such as the skin and muscles. Likewise, the increased spontaneous activities of TG neurons should include the activities of afferents projecting to the TMJ and adjacent orofacial tissues. The VF assay was performed on the skin overlying the TMJ, and we acknowledge that nocifensive responses by VF cannot be considered clinical pain in response to pressure on the TMJ. The low range of EF_50_ in mice has indicated that the nocifensive responses are more likely due to the activation of cutaneous rather than deep tissue nociceptors. As increased pressure pain is the most frequent symptom of patients with TMJ arthralgia [37], methods to assess pressure pain directly on the TMJ in mice need to be developed in the future. Similarly, in in vivo Ca^2+^ imaging, mechanical, thermal, and chemical stimuli were delivered to the skin in the mandibular area; therefore, the activated TG neurons should be primarily cutaneous afferents. Therefore, our results suggest that FMO-induced TMJ injury leads to robust peripheral sensitization of the cutaneous and TMJ afferents. In patients with TMJ arthralgia, pressure pain and mechanical and cold hyperalgesia on the skin overlying the TMJ are evident [15; 37]. However, the lack of thermal hypersensitivity after FMO in mice was inconsistent with thermal hyperalgesia on the skin overlying the TMJ in patients with TMJ arthralgia [15; 37]. We do not presume that all the observed behavioral hyperalgesia can be exclusively attributable to peripheral mechanisms. Although bite force reduction induced by masseter inflammation is attributable to the peripheral mechanisms in mice [36], technical limitation of in vivo Ca^2+^ imaging does not allow us to determine the association of peripheral sensitization and bite force reduction following TMJ injury. Increased failure of successful biting after FMO likely reflects the increased emotional affective pain mediated by plastic changes across higher order brain structures following TMJ injury, rather than direct consequences of peripheral sensitization. Likewise, anxiety-like behaviors are more likely due to the consequences of central sensitization. Although we did not determine the extent of central sensitization after FMO, previous studies showed that similar TMJ injury can produce central sensitization. Cytokines and neuropeptides are upregulated in the upper spinal cord and trigeminal nucleus caudalis [9; 30]. Prolonged mouth opening in rats leads to changes in brain networks, including prefrontal-limbic pathways, which is likely associated with persistent pain [29]. Nevertheless, central sensitization is initiated and maintained by peripheral inputs and peripheral sensitization can play critical roles.

Our study suggests there are several different mechanisms of peripheral sensitization following TMJ injury. FMO-induced TMJ injury leads to robust inflammatory responses in TMJ. It increases inflammatory cytokines such as TNF [13; 32], which likely induce sensitization of the peripheral terminals of TMJ afferents. Soluble factors or immune cells can spread from the TMJ to the adjacent skin to induce the sensitization of cutaneous afferents. In addition to the canonical mechanisms of peripheral sensitization, our in vivo Ca^2+^ imaging study showed increased spontaneous activities of TG neurons not only in the V3 area but also across the V1/V2 areas of the TG. This finding suggests intra-ganglionic spreading of peripheral sensitization, which can mediate ectopic inflammatory and neuropathic orofacial pain [1; 26]. Our histological analysis also indicated another potential mechanism of spreading hyperalgesia following TMJ injury. We observed a significantly greater number of CD45+ immune cells within the nerve bundle around the TMJ in the FMO group than in the sham group, indicating intraneural inflammation following injury. Chronic constriction injuries induce intraneural inflammation that exacerbates axonal loss and neuropathic pain [8]. Intraneural DAPI+ cells were also substantially greater after FMO, suggesting that glial cells, such as Schwann cells, in nerve bundles are increased following FMO. The pronociceptive role of Schwann cells in neuropathic pain is well known [38], and the inflammatory proliferation of glial cells in peripheral nerves may lead to hyperalgesia. In humans, the TMJ is innervated primarily by the two branches of the mandibular nerve (Fig. 10F) [16; 22]. The auriculotemporal nerve is a branch that innervates the posterior part of the TMJ capsule and retrodiscal tissues, then takes a lateral course and innervates the periauricular skin. The masseteric nerve branch proceeds laterally, innervates the anterior capsule of the TMJ, passes over the lateral pterygoid muscle, and eventually innervates the masseter muscle. Therefore, inflammation in the TMJ can potentially sensitize the nerve bundles passing through it, innervating adjacent orofacial structures. Consequently, intra-articular inflammation could cause hyperalgesia in the TMJ and contribute to hyperalgesia spreading to adjacent structures through peripheral sensitization via intraneural inflammation. Further study is warranted on the potentially novel and unique peripheral mechanisms of spreading hyperalgesia in the TMJ.

Our mouse model of TMJ injury also has limitations. The FMO model shows no clear sex differences in behavioral hyperalgesia and primary afferent hypersensitivity, which is not consistent with the higher prevalence of TMDs in women than men [2; 19]. The time course of the behavioral phenotypes and associated peripheral sensitization is not long enough to mimic the clinical manifestation of chronic pain conditions. The association of the model with psychological status is unclear either. Nonetheless, the FMO mouse model can be a clinically relevant translational model for studying post-traumatic pain-like behaviors over subacute time course for determining the underlying peripheral mechanisms of TMJ pain.

## Acknowledgments

The authors thank Dr. Ke Ren and Dr. Ken Hargreaves for critical reading of the manuscript. This study was supported by National Institutes of Health Grant R01DE026677 (YSK), R01NS128574 (YSK), R01DE031477 (YSK and MKC), R35DE030045 (MKC), R01 DE016062 (JYR), University of Maryland School of Dentistry IN-SPIRE grant program (MKC), and T32 DE014318 University of Texas Health Science Center Craniofacial Oral-biology Student Training in Academic Research (COSTAR) (JS). IA receives a fellowship from King Saudi Arabia.

